# Complementary effects of above- and belowground biodiversity on ecosystem functions across global grasslands

**DOI:** 10.1101/2023.02.10.528097

**Authors:** Catarina S. C. Martins, Manuel Delgado-Baquerizo, Ramesha H. Jayaramaiah, Jun-Tao Wang, Tadeo Sáez-Sandino, Hongwei Liu, Fernando T. Maestre, Peter B. Reich, Brajesh K. Singh

**Affiliations:** Hawkesbury Institute for the Environment, Western Sydney University, Penrith, NSW, Australia; Laboratorio de Biodiversidad y Funcionamiento Ecosistémico. Instituto de Recursos Naturales y Agrobiología de Sevilla (IRNAS), CSIC, Av. Reina Mercedes 10, E-41012, Sevilla, Spain; Unidad Asociada CSIC-UPO (BioFun). Universidad Pablo de Olavide, 41013 Sevilla, Spain; Departamento de Ecología, Universidad de Alicante, San Vicente del Raspeig, Alicante, Spain; Instituto Multidisciplinar para el Estudio del Medio “Ramón Margalef”, Universidad de Alicante, San Vicente del Raspeig, Alicante, Spain; Department of Forest Resources, University of Minnesota, Saint Paul, MN, USA; Institute for Global Change Biology, School for Environment and Sustainability, University of Michigan, Ann Arbor, Michigan, USA; Global Centre for Land-Based Innovation, Western Sydney University, Penrith, NSW, Australia

## Abstract

Grasslands are integral to maintaining biodiversity and key ecosystem services under climate change. Plant and soil biodiversity, and their interactions, support the provision of multiple ecosystem functions (multifunctionality). However, whether plant and soil biodiversity explain unique, or shared, contributions to supporting multifunctionality across global grasslands remains virtually unknown. Here, we combine results from a global survey of 101 grasslands with a novel microcosm study, controlling for both plant and soil microbial diversity to identify their individual and interactive contribution to support multifunctionality under aridity and experimental drought. We found that, plant and soil microbial diversity independently predict a unique portion of variation in above- and belowground functioning, suggesting both types of biodiversity complement each other. Interactions between plant and soil microbial diversity regulated primary productivity, nutrient storage, and plant productivity. Our findings were also context dependent, since soil fungal diversity was strongly associated to multifunctionality in less arid regions, while plant diversity was strongly linked to multifunctionality in more arid regions. Our results highlight the need to conserve both above- and belowground diversity to sustain grassland multifunctionality in a drier world and indicate climate change may shift the relative contribution of plant and soil biodiversity to multifunctionality across global grasslands.

## Introduction

Grasslands are one of the major biomes of the world, covering about 40% of the Earth’s surface and 69% of the Earth’s agricultural land area (Suttie et al., 2005; O’Mara, 2012; Bengtsson et al., 2019). These ecosystems are globally recognized for their high biodiversity (Habel et al., 2013; Bond, 2016) and are also essential for the provisioning of a wide range of ecosystem services, including food production (O’Mara, 2012), carbon (C) storage and climate mitigation (Yang et al., 2019), pollination (Sexton & Emery, 2020), water regulation and a range of cultural services (Bardgett et al., 2021). Despite their importance, grasslands are increasingly under threat of degradation at an alarming pace in many parts of the world due to anthropogenic activities including climate change (Huang et al., 2016; Burrell et al., 2020; Bardgett et al., 2021). Part of this degradation is associated with the loss of their biodiversity. Furthermore, above- and belowground biodiversity are tightly linked, with plants providing energy belowground *via* plant litter and root exudates, and soil microorganisms supporting nutrient availability for plants through fixation and organic matter (OM) decomposition (Hooper et al., 2000; Wardle et al., 2004; Van Der Heijden et al., 2008). However, there is little understanding on how plant and soil biodiversity interact under environmental stresses to regulate the provision of multiple ecosystem functions (multifunctionality) in grasslands across contrasting environmental conditions.

The positive relationship between plant diversity and ecosystem functioning has been a focus of research for more than two decades (Tilman et al., 1996; Isbell et al., 2011; Maestre et al., 2012; Reich et al., 2012; Tilman et al., 2014), with similar findings recently expanded to belowground communities in terrestrial ecosystems across biomes (Jing et al., 2015; Delgado-Baquerizo et al., 2016). In both cases, the higher the plant or soil biodiversity, the higher the ecosystem multifunctionality supported by terrestrial ecosystems. Further, the effects of plant and microbial diversity are largely expected to be complementary (Barry et al., 2019), maximizing rates of key above- and belowground services (Bardgett & Wardle, 2010). However, our current knowledge and lack of strong experimental evidence does not allow us to tease apart the relative contribution of, and interactions between plant and soil microbial diversity in driving multifunctionality, as the influence of plant and soil biodiversity are often evaluated in isolation. Moreover, recent studies suggest that plant and soil biodiversity rarely peak in the same locations across the globe (Cameron et al., 2019; Delgado-Baquerizo et al., 2019), highlighting an urgent need for quantification of the relative contribution of plant and soil biodiversity to support grasslands multifunctionality.

Current uncertainties in the relative contribution of plant and soil microbial diversity in regulating multifunctionality in a climate change context exist because of three main reasons. First, the interaction between plant and soil biodiversity is often overlooked in global studies and manipulative experiments (Wagg et al., 2014; Bardgett et al., 2014; Delgado-Baquerizo et al., 2016, 2020). However, this knowledge is critical to understand whether these two groups can work independently and/or synergistically in supporting ecosystem functions. For example, soils with high plant and microbial diversity may support higher rates of nutrient cycling and decomposition compared to soils with either low plant or microbial diversity (Cameron et al., 2019) but strong empirical evidence is lacking. Second, most studies investigating grasslands multifunctionality are based on local and regional studies and lack experimental support manipulating plant and soil biodiversity simultaneously (Jing et al., 2015; Soliveres et al., 2016; Hu et al., 2021). Global field survey observations are required to establish biodiversity-ecosystem functioning relationships and to understand the relative contribution of above- and belowground diversity in a real-world scenario, and how it is influenced by key abiotic (*e.g.,* soil types) and climatic (*e.g.,* aridity) conditions. Moreover, mechanistic evidence from manipulative experiments that simultaneously quantify the influence of plant and soil microbial diversity in driving ecosystem multifunctionality are needed to distinguish between statistical correlation and causal relationships. However, manipulative experiments that simultaneously account for plant and microbial diversity were not feasible until recently, mainly due to a technical inability to maintain soil biodiversity gradients over a sustained period in the presence of plant communities (Yang et al., 2021). Finally, current rate of biodiversity loss is associated with significant change in climatic conditions but there is lack of experimental evidence on how climate change variables impact the relationship between plant and soil microbial diversity and ecosystem multifunctionality in the context of on-going global environmental changes. Addressing these knowledge gaps is critical to advance our understanding of plant-soil feedback effects, predict the consequences of current environmental disturbances and to develop effective management and conservation policies to maintain and restore global grasslands.

Here, we aimed to (1) identify and quantify the unique and interactive contribution of microbial and plant diversity in driving multiple ecosystem functions in global grasslands; and (2) assess the influence of observational (aridity) and experimental (drought) climatic conditions on the linkages between biodiversity and functions. We chose aridity and drought treatments because both represent changes in water availability and potentially illicit similar community responses. We hypothesized that plant and soil microbial richness explain a unique portion of variation in the distribution of ecosystem multifunctionality across global grasslands under climate change, supporting the argument that plant and soil biodiversity have a complementary effect on ecosystem functions (Figure 1a). To achieve this, we combined a global field survey including plant and soil biodiversity and multiple functions in 101 grasslands across biomes (from arid to tropical grasslands; Figure 1b) with a novel manipulative experiment of plant and soil biodiversity using four levels of plant richness and three levels of soil microbial richness in a full-factorial design subjected to drought stress (Figure 1c). The main aim of our manipulative experiment was to determine the direct effects and relative contribution of plant and soil biodiversity in explaining multifunctionality under contrasting water availability conditions. Soil microbial diversity-ecosystem functioning studies suffer from major limitations, particularly survey-based studies that link natural variation in microbial communities to change in ecosystem functions. However, disentangling cause and effect from observational (survey-based) studies is difficult as other drivers (*e.g.* temperature, soil structure) could simultaneously impact both community and ecosystem functions (Crawford et al., 2012). To account for these, our experimental framework for the manipulative experiment, used one soil type to avoid cofounding impacts of soil characteristics such as structure and nutrients availability. By also using a dilution to extinction approach, all parameters of soils remained the same except for a change in soil biodiversity, allowing the identification of direct impacts of soil biodiversity on ecosystem functions (Trivedi et al., 2019; Delgado-Baquerizo et al., 2020). Our unique approach combining two independent methods (observational and experimental) provides a complementary assessment of the linkages between plant and soil microbial diversity and ecosystem multifunctionality. Our results identify, for the first time, the complementary roles of plant and soil biodiversity in explaining multiple aspects of functions in grasslands under experimental climate change, and across environmental gradients.

**Figure 1.**
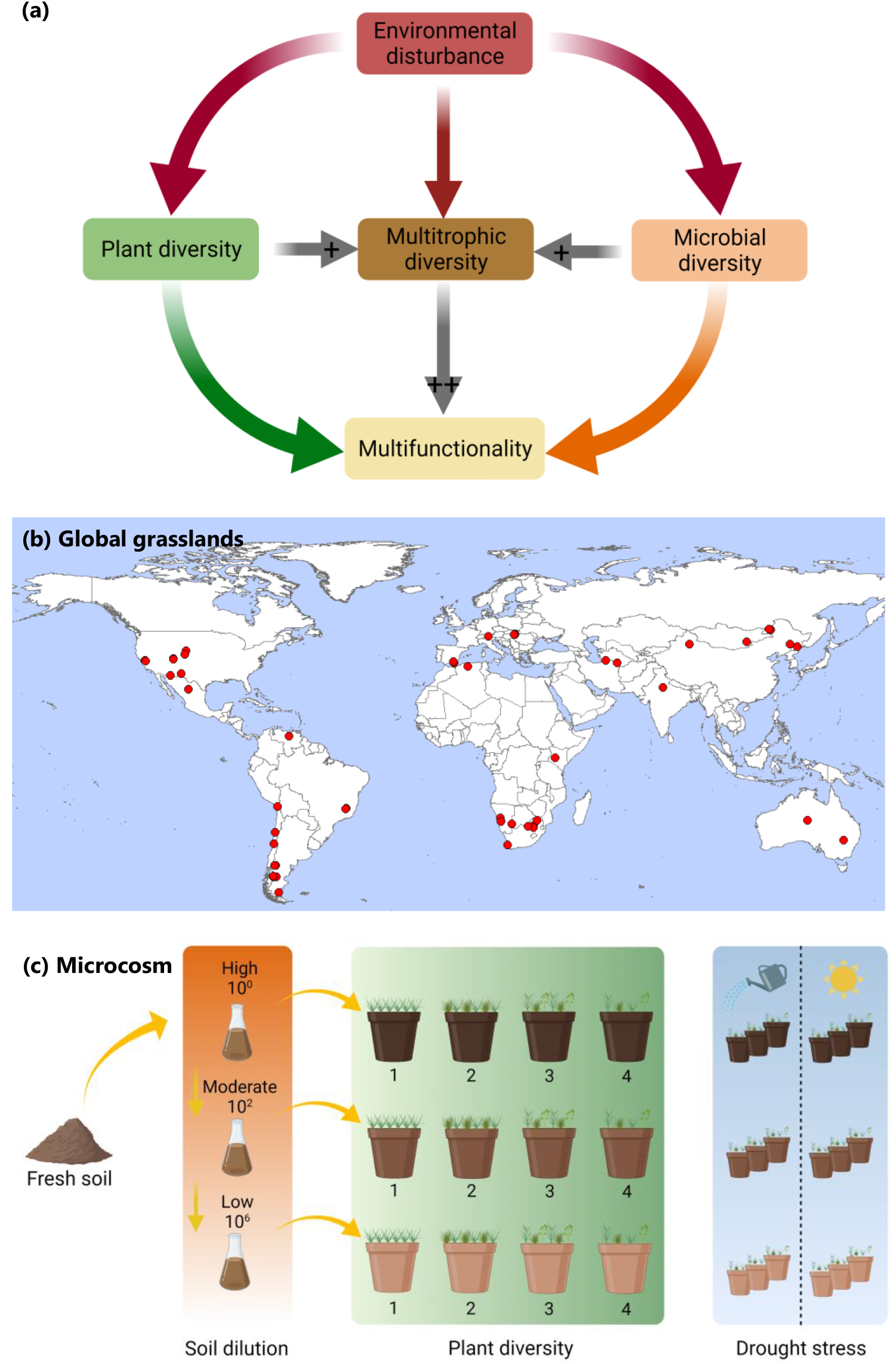
Experimental framework to study the effects of plant and microbial diversity interactions on ecosystem multifunctionality. (a) conceptual framework exploring the relative contribution of plant and microbial diversity and their combination, *i.e.,* multitrophic diversity to multiple ecosystem functions; (b) locations of the 101 sites included in the global grassland survey; (c) full-factorial design of the microcosm study comprising four plant diversity levels of plant species and three microbial diversity levels obtained from a dilution-to-extinction approach; (created with BioRender.com).

## Results

### Plant and soil microbial richness are linked to ecosystem multifunctionality across global grasslands

To investigate the relationships between above- and belowground biodiversity and multifunctionality, we first used regression analysis and Spearman correlations. In soils from globally distributed grasslands, we found significant positive relationships between plant, bacterial, and fungal richness and the combination of plant and microbial richness, *i.e.,* multitrophic richness, with weighted multifunctionality (Figure S7a-c). We used weighted multifunctionality (a composite, standardized metric, see Methods) to account for the equal contribution of the three groups of ecosystem functions included in the global survey, but also to improve results comparison from the two independent studies. Averaged multifunctionality was also tested for comparison (Figure S7b). From here on, weighted multifunctionality was used and referred to as multifunctionality. We found strong and positive correlations between multifunctionality and fungal richness, particularly mycorrhizal and saprotrophic fungi, followed in strength by multitrophic richness (Figure 2a). Furthermore, relations with multifunctionality were strongest for plant richness in arid and hyper-arid environments, for fungal richness (including mycorrhizal and saprotrophic fungi) in humid and arid environments, and for bacterial richness in dry sub-humid and semi-arid environments (Figure S8). Our results also showed that plant and soil microbial richness explained unique portions of variation in multifunctionality, which were not explained by environmental properties alone, using a standardized multi-model inference approach (Figure 2b). Specifically, environmental properties explained 46% of the variance (with the inclusion of aridity (5%)) in comparison to the richness of plants (29%) and soil microbes (26%) (Figure 2b).

**Figure 2.**
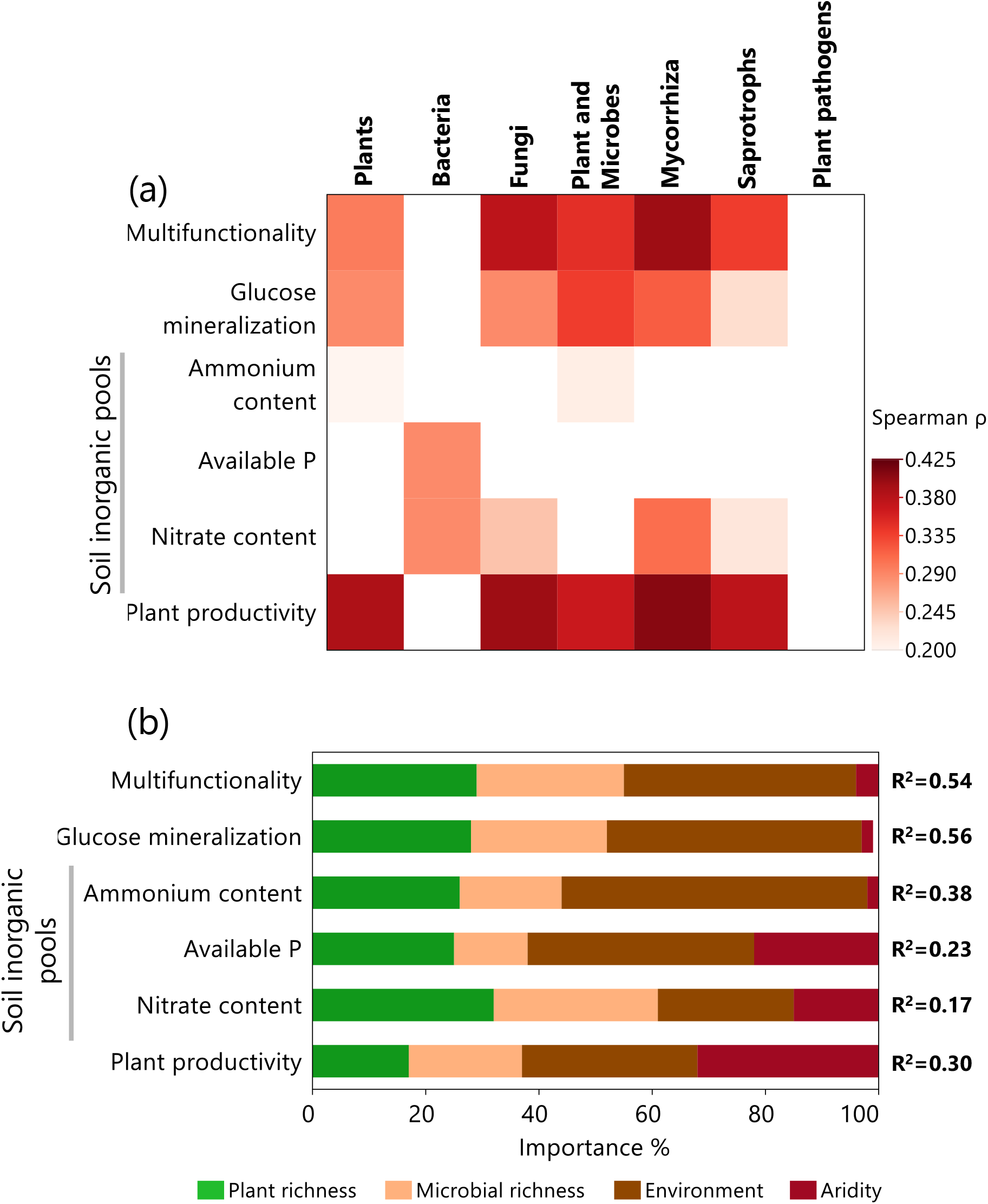
Links between ecosystem multifunctionality and plant and soil microbial richness in a grassland field survey. (a) Significant correlations (Spearman; *p* ≤ 0.05) between the richness of groups of organisms and weighted multifunctionality and individual ecosystem functions. (b) The contribution of plant and microbial (bacteria and fungi) richness, environment (distance from the equator, plant cover, soil pH, % clay, soil C and mean annual temperature) and aridity index to weighted multifunctionality and individual functions (glucose mineralization – soil respiration with glucose addition; soil inorganic pools – ammonium content, available P, nitrate content; plant productivity – net primary production). Microbial richness corresponds to a composite metric of their joint richness (standardized between 0 and 1) and environment properties correspond to a standardized principal component analysis first axis obtained from the multiple properties. The importance of predictors is expressed as the percentage of explained variance, taken as the absolute value of their standardized regression coefficients. R^2^ values express total variances corresponding to model adj. R² obtained from parameter estimates averaged across all models. This method is similar to a variance partition analysis because predictors have been standardized.

Structural equation modelling analysis further illustrated the direct impact of microbial richness on multifunctionality, after accounting for environmental properties of spatial influence, climate, soil and plant variables (Figure 3a). Plant richness had indirect effects on multifunctionality *via* direct effects on soil bacterial and fungal richness. Our analyses also showed that the mean ambient temperature (MAT) indirectly impacted multifunctionality *via* plant richness and microbial richness. In contrast, the impacts of aridity on multifunctionality were indirect *via* direct and negative effects on soil characteristics such as soil C and % of clay (the higher the aridity index the lower the aridity; Figure 3a).

**Figure 3.**
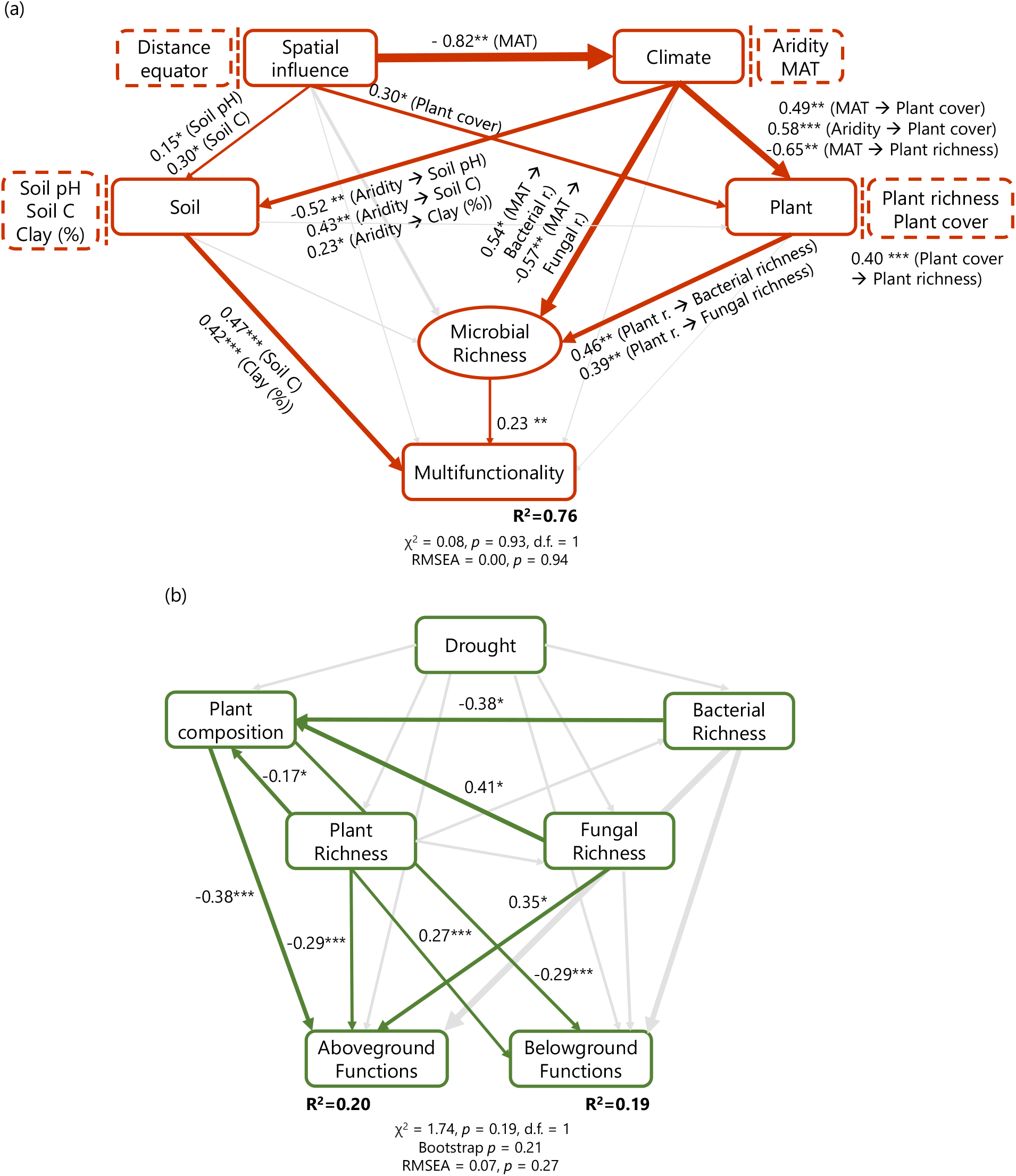
Structural equation models for (a) weighted multifunctionality derived from the global grassland survey (n=101) and for (b) above- and belowground functions derived from the microcosm study (n=157). We aimed to identify the direct relationship between plant and soil biodiversity (bacterial and fungal richness) and multiple ecosystem functions. In the case of the global survey, a similar approach to Delgado-Baquerizo et al., (2020) was used, by grouping different categories of predictors (climate, soil properties, plants and spatial influence) into the same box in the model for graphical simplicity; however, these boxes do not represent latent variables. Soil microbial richness was included as a composite variable, including information of soil bacterial and fungal taxa. Please note, the higher the aridity index the less arid environment it represents. The rectangles are observable variables. In the case of the microcosm study, plant composition corresponds to first axis of a principal component analysis. Numbers adjacent to arrows are indicative of the effect size of the relationship. For each model, the proportion of variance explained (R^2^) and the various goodness-of-fit statistics are shown below the response variables. Significance levels are as follows: \**p* ≤ 0.05, \*\**p* ≤ 0.01 and \*\*\**p* ≤ 0.001. Abbreviations: MAT, mean annual temperature; RMSEA, root mean square error of approximation. Information about a priori models can be found Table S5a and S5b.

We conducted additional analyses to evaluate the effect of biodiversity on the performance of multifunctionality when considering an increasing number of functions (multi-threshold multifunctionality approach; see methods). We observed that mycorrhizal and saprotrophic fungal richness had a significant and positive association with functions at low (10%, 25%) and medium (50%) thresholds of performance, *i.e.*, significant association above the threshold (%) of their maximum observed levels of rate/availabilities (Figure S9a). Multitrophic richness was only positively related to functions at low thresholds of over 10% and 25%, followed by plant richness associated to functions of only over 10% of their maximum observed levels of functioning (Figure S9a). These results indicate that soil and plant diversity are both necessary to support a wide range of rate/availabilities of multiple functions (*i.e.*, functions performing at only 10% of their maximum), supporting a fundamental level of functioning in grasslands.

### Assessing the relative contribution of plant and soil microbial richness as multifunctionality drivers in a microcosm study

Linear mixed modelling showed that plant and soil microbial richness had independent effects on multifunctionality in the experimental microcosm study (Table S6a-b; Figure S6). Experimental losses in microbial richness had negative effects on both above- and belowground functions, such as leaf nitrogen (N) content, plant productivity (green canopy cover and plant height), soil nutrient storage (soil C, N and, phosphorus (P)), and soil inorganic pools (inorganic N), whilst it increased (positive effect) only leaf P content and soil dissolved organic C (DOC). Increasing plant richness decreased leaf C content, whereas it had positive effects on belowground functions such as total dissolved N (TDN), soil nutrient storage (soil P) and OM decomposition (glucose mineralization). The drought event significantly increased soil nutrient storage (total N and total P) and consequently, multifunctionality.

Regression analysis and Spearman correlations showed a significant positive relationship between multifunctionality (obtained from six grouped functions) and plant and multitrophic richness (Figure 4a; S7d-f). Multitrophic richness was responsible for maintaining a positive relationship with belowground services, specifically soil nutrient storage and OM decomposition (Figure 4a). In particular, we found a positive relationship between plant richness and soil N stocks (TDN and total N), and OM decomposition (basal respiration and glucose mineralization), whilst fungal richness was positively and negatively related to C stocks (total C) and labile C (DOC), respectively (Figure 4a). We then tested the biodiversity-multi-threshold multifunctionality relationship and found a positive association between plant and multitrophic richness and functions at high thresholds of over 50% and 90% of their maximum observed levels of functioning, respectively (Figure S9b).

**Figure 4.**
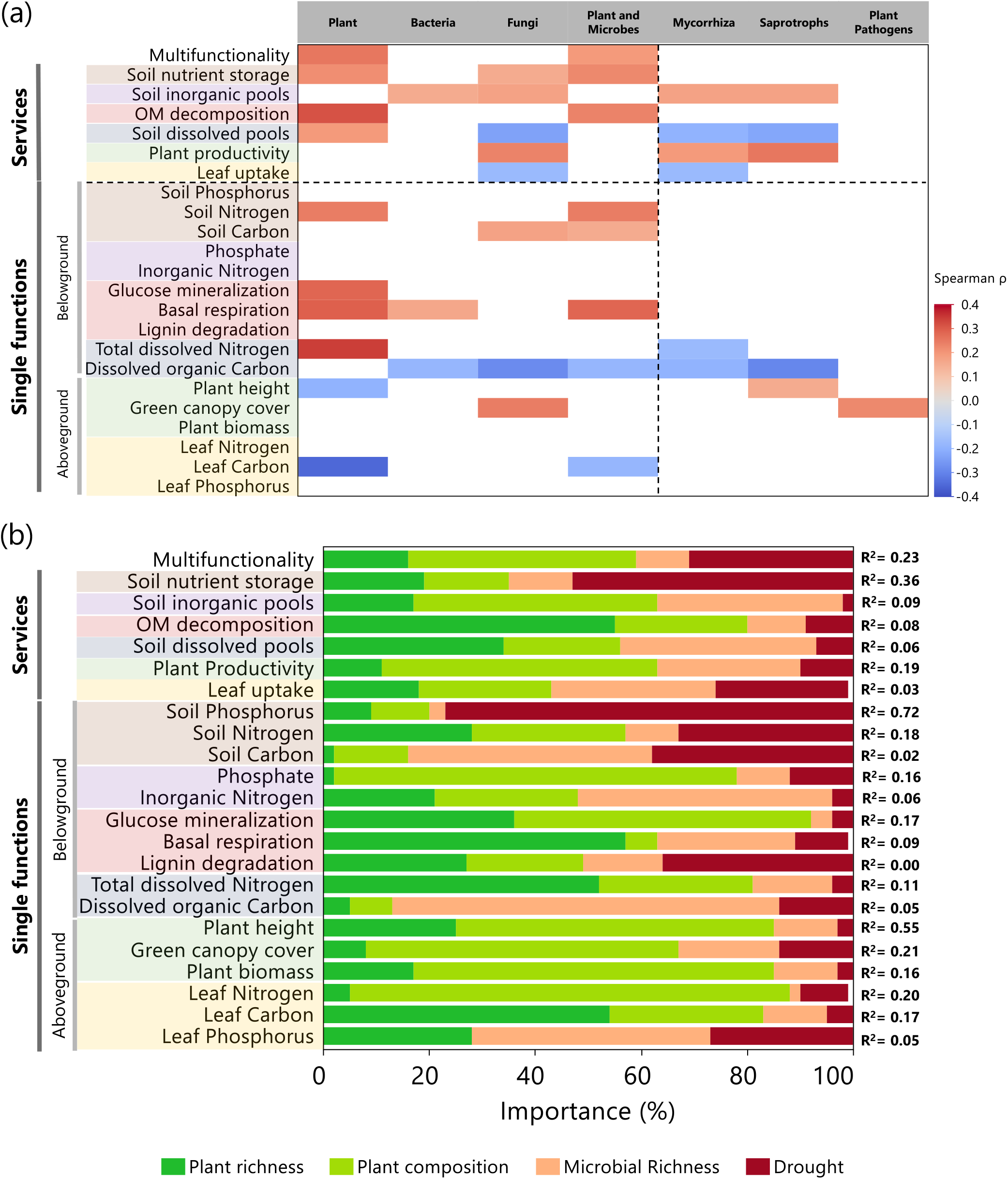
The contribution and relationship of plant and soil biodiversity to weighted multifunctionality. (a) Significant correlations (Spearman; *p* ≤ 0.05) between richness groups and weighted multifunctionality, services and individual belowground and aboveground functions. The different background colours are used to highlight which individual functions belong to services categories. Plant and microbes richness corresponds to a composite metric of their joint diversity (standardized between 0 and 1). Fungal phylotypes include mycorrhiza, saprotrophs and plant pathogens fungi. (b) The contribution of plant and microbial (bacteria and fungi) richness, plant composition and drought to weighted multifunctionality, services and single aboveground and belowground functions. The different background colours are used to highlight which single functions belong to services categories. The importance of predictors is expressed as the percentage of explained variance, taken as the absolute value of their standardized regression coefficients. R^2^ values express total variances corresponding to model adj. R² obtained from parameter estimates averaged across all models. This method is similar to a variance partition analysis because predictors have been standardized. Microbial richness encompass total bacteria and total fungi.

To test the influence of community composition and abundance on the relationship between richness and multifunctionality, we used partial correlations and found that the positive effects of plant richness on multifunctionality were marginally affected by plant composition and abundance. The lack of significant effects of bacterial and fungal richness on multifunctionality remained unchanged when controlling for microbial abundance and community composition (Table S7). Thus, we included plant composition (*i.e.*, different plant combinations), together with plant and microbial richness and drought to assess their importance and contribution as predictors of multifunctionality, and individual functions. We found that plant (16%) and soil microbial (10%) richness predicted a combined unique portion of variation (26%) on a similar order of magnitude with plant composition (43%) and drought disturbance (31%). The percentage of explained variance driven by drought was on average higher for belowground functions with 23 ± 7 %, in comparison to 11 ± 4 % for aboveground functions (Figure 4b), with strongest impacts on soil nutrient storage, particularly 77% for soil P (Figure 4b). Our findings also suggest fungal richness may have been indirectly influencing plant composition importance in predicting aboveground functions (Figure 4b; Table S6b) as supported by structural equation modelling analysis (Figure 3b).

## Discussion

By combining a global grassland survey and a unique manipulative microcosm experiment, our study showed that: (1) plant and soil microbial diversity independently predict a unique portion of variation in above- and belowground ecosystem functioning, suggesting that both plant and soil biodiversity complement to support functions across global grasslands subjected to climate change (aridity gradients and drought) and; (2) multitrophic biodiversity, *i.e.,* plant and soil microbial richness combined, is positively associated with multifunctionality, and was essential to maintain key ecosystem services such as soil nutrient storage and primary production globally. This highlights the multiple pathways in which plant and soils can connect to support soil fertility and nutrient cycling but also plant productivity.

We further illustrated that at a global level, grassland multifunctionality varied from a stronger positive association with fungal richness in humid regions, to a stronger positive association with plant richness in hyperarid regions, whilst in arid regions both plant and fungal richness were strongly associated with multifunctionality. In addition, aridity indirectly explained global multifunctionality by a negative association with soil C and % of clay, while the drought event implemented experimentally only temporarily impacted multifunctionality in the microcosm study, which substantially recovered. Instead, the experimental drought event led to an increase in soil nutrient storage, particularly soil P concentration in the soil (Delgado-Baquerizo et al., 2013), likely caused by temporary reduced microbial demand and nutrient uptake by plants in response to drought event (Manzoni et al., 2012; Schimel, 2018). These findings support the necessity in maintaining key ecosystem functions through maintenance of biodiversity in vulnerable ecosystems which are projected to face increasing water scarcity under climate change conditions.

Our global survey showed that the richness of soil fungal groups, particularly mycorrhizal and saprotrophic fungi, were strongly and positively associated to ecosystem multifunctionality across grasslands. Mycorrhizal fungi are known to promote a range of benefits to plants by providing mineral nutrients and protection from abiotic and biotic stresses (Chen et al., 2018; Begum et al., 2019; Tedersoo et al., 2020). Saprotrophic fungi are essentially soil decomposers capable of breaking down organic matter otherwise unavailable to plant growth (Gessner et al., 2010), due to their extensive hyphal networks (de Vries et al., 2018) and their capacity to breakdown more recalcitrant forms of organic matter (Eastwood et al., 2011). Thus, fungal communities provide critical support for both above- and belowground communities and functions. Additionally, the positive relationship between multitrophic richness and multifunctionality was weaker in comparison to individual richness groups. In fact, positive effects of plant richness on multifunctionality at a global level were a result of its indirect effects on microbial richness, as previously observed across biomes (e.g., forests, grasslands and shrublands) (Delgado-Baquerizo et al., 2016; 2020), suggesting plant and microbial richness have selective effects when driving specific ecosystem services. For example, our microcosm study demonstrated a strong, positive, and significant association of multitrophic richness with soil C and N storage and OM decomposition, whereas the importance of plant productivity was highlighted in the global survey. This is indicated in the microcosm study by the promotion of soil N stocks by plant richness which may have been attributed to N inputs being made available by increasing microbial activity and more labile forms of OM decomposition. In contrast, fungal richness promoted C stocks and consumed labile C (DOC), suggesting the higher C storage observed in soils with high microbial richness was likely driven by fungal retention, possibly from photosynthates derived from higher green canopy cover (Orwin et al., 2011).

Our results also suggested that the strongest positive association was observed between fungal richness and multifunctionality at low and medium thresholds of <50% across global grasslands. While in the microcosm study, an association between plant richness and functions operating at medium and high levels of functioning (>50% threshold) was observed. These results indicate that fungal and plant diversity are both necessary to sustain a wide range of rate/availabilities of multiple functions, supporting a fundamental level of functioning in grasslands. In particular plants, as primary producers, directly affect resource availability and biomass production (Cardinale et al., 2011), and thus are expected to have a significant impact in the provisioning and stability of multiple services (Jing et al., 2015; Soliveres et al., 2016). This is also supported by the fact that plant composition in the microcosm study explained a unique variation of multifunctionality demonstrating the types and abundance of plant species also strongly influence ecosystem functions (Carrick & Forsythe, 2020; Bowker et al., 2021).

Our study provides new insights on the fundamental importance of plant and soil biodiversity to support grassland multifunctionality in a context of climate change, and water availability in particular. Both our datasets illustrate that the impacts of reduced water availability, *i.e.,* aridity in global natural grasslands and a single experimental drought event in the microcosm study, influence certain aspects of ecosystem multifunctionality. However, our findings also suggest that globally, the decline in soil fertility under increasing aridity and increasing temperatures will markedly influence multifunctionality. Thus, indirect effects of aridity are likely to have major effects on multifunctionality. For example, the decline of C and clay content that we observed with aridity could be associated with the decline in plant productivity and plant cover and the likely decrease in plant-derived organic inputs into the soil (Berdugo et al., 2020). Additionally, the sharp decline observed in the fungal richness, namely saprotrophic and mycorrhizal fungi, from hyper-arid regions could be a response to high mean annual temperatures and the decline of vegetation. In fact, Berdugo et al. (2020) refers to progressive decline phases in ecosystem functioning as a response to aridity until an irreversible phase is reached. Vegetation is expected to be impacted first, followed by soil fertility, and only then microbial communities such as saprotrophic fungi, known to be key drivers of soil fertility (Ochoa-Hueso et al., 2018), and mycorrhizal fungi known to be linked to plant composition and nutrient cycling (Lu & Hedin, 2019). Our experimental study illustrated an almost complete recovery of multifunctionality, likely because it only included a single extreme drought event. More frequent drought disturbances would probably have more permanent and deleterious effects in both above- and belowground functions, constraining biodiversity-multifunctionality relationships to recover. We expectedly found some difference in results such as the relative contributions of microbial diversity on multifunctionality between survey and experimental studies given restrictions of space for root foraging and microbial dispersal in pots. However, key findings overlap supporting our hypothesis. Overall, we provide novel empirical evidence that maintaining both plant and microbial diversity is crucial to sustain multiple ecosystem functions due to their independent and interactive roles in ecosystem functioning under climate change. In particular, we showed that plant and soil biodiversity explained a unique portion of variation in the distribution of multifunctionality across global grasslands and in experiments subjected to climatic stress. These findings support that plant and soil biodiversity complement each other to support functioning under environmental stress. Fungal richness had a direct impact on multifunctionality in global grasslands whilst plant richness effects were indirectly driven by microbial richness, particularly for aboveground functions. Multitrophic richness was vital in maintaining belowground functions, particularly soil nutrient storage and OM decomposition but also plant productivity globally. We provide novel insights that fungal richness has a stronger positive association with multifunctionality in humid regions whilst plant richness in hyper-arid regions and both in arid regions, indicating context dependency in biodiversity and ecosystem functions relationships. Overall, our study provides empirical evidence and supports that both above- and belowground diversity and their interactions in terrestrial ecosystems should be explicitly considered in future ecosystem biodiversity and functioning studies and in developing new management, restoration, and conservation policies for grasslands.

## Material and methods

### Global survey

#### Study sites

We used composite topsoil samples from global field surveys, which were conducted between 2014-2019 following standardized field protocols. This global field survey includes 101 grasslands from six continents, providing a large representation of all climatic grassland biomes on the planet, *i.e.*, humid, dry-subhumid, semi-arid, arid and hyperarid (Figure 1b). Grasslands were dominated by globally dominant grass genera such as *Festuca*, *Stipa*, *Andropogon*, *Bouteloua*, *Cynodon*, *Eragrostis*, *Poa*, *Sporobolus* and *Trachypogon*. These grasslands also included a wide range of environmental conditions such as soil pH (4.3-8.6), fine texture (1.7-83.6%), carbon (0.4-215.1 g C kg soil^-1^), mean annual precipitation (26-1471 mm) and temperature (-2.7 – 27.2°C). Perennial plant richness and plant cover were determined in the field using the line-intercept method according to Maestre et al. (2012).

Composite soil (top ∼10 cm) samples (from 5-10 soil cores) were collected in these locations following the protocol described in Maestre et al. (2012) aiming to capture the within-location heterogeneity variation in soil environmental conditions. A portion of these soils was frozen (– 20 °C) after sampling for molecular analyses, while another portion was air-dried and used for determining soil properties.

#### Microbial diversity characterization

Bioinformatic analysis on the 16S rDNA/ITS amplicons was performed as below. Briefly, pair-end reads were merged using USEARCH (Edgar, 2010), followed by a quality filtering step on the merged reads with expected error (ee) set as 1.0 maximum. High quality reads were then dereplicated and singletons were removed. The zOTUs (denoised sequences, 100% sequence identity) were gained by denoising (error-correction) the amplicon reads using unoise3 (Edgar & Flyvbjerg, 2015). Representative sequences of 16S rDNA and ITS zOTUs were annotated against the Silva (Quast et al., 2013) and UNITE (Nilsson et al., 2018) database in QIIME (Caporaso et al., 2010) using the UCLUST algorithm (Edgar, 2010), respectively. Approximately 15.8M and 17.1 high-quality merged-sequences were mapped for 16S rDNA and ITS zOTUs, respectively. Prior to diversity calculation, a normalization procedure was performed at 15,000 and 11,000 sequences *per* sample for 16S rDNA and ITS tables, respectively. Alpha diversity indices, including richness and Shannon diversity, and beta diversity indices including Bray-Curtis matrices were calculated in QIIME. The FUNGuild database was employed to gain the functional guilds information of fungi (Nguyen et al., 2016).

#### Ecosystem Functions

We selected five ecosystem functions from those available in the global survey to match those functions available from the microcosm study. These five functions were further grouped in three ecosystem services: OM decomposition (glucose induced respiration), soil inorganic pools (availability of nitrate, ammonium and phosphate), and plant productivity (net primary productivity; NPP). The five functions/services considered are mostly weakly correlated with each other (Table S4b), supporting their inclusion in the multifunctionality index. We used NDVI (Normalized Difference Vegetation Index), from MODIS satellite imagery (https://modis.gsfc.nasa.gov) (2001-2020; 250m resolution), as our proxy of NPP. The content of nitrate and ammonium were determined as a proxy of nitrogen availability as described in Maestre et al. (2012). The availability of these two N forms is associated with important processes such as nitrification and ammonification rates. The content of soil phosphate was determined using Olsen P approach as described in Maestre et al. (2012). The respiration of soil in response to glucose was determined at 20°C and 60% WHC using Microresp® as described in Campbell et al. (2003).

### Microcosm: experimental design

The microcosm study was conducted at Western Sydney University (WSU) for approximately six months in a full-factorial design, including four levels of plant richness, three levels of microbial diversity, a drought event at the end (well-watered control vs. drought) and six replicates for each combination of treatments. Soils were collected from the experimental grassland research facility at WSU. Briefly, each plant species (originally six; see plant diversity manipulation) was represented in monoculture, in three different three species combinations, and one final combination of six species (1, 3, 6 species). In addition, 18 control replicates with no plants and with only the microbial diversity treatment were established, totalling 198 microcosms. Plant richness was monitored throughout the experimental period: a preliminary assessment was made in the first 12-weeks of plant establishment (T1); after 6-weeks of plant establishment and before drought was initiated (T2); at the end of the 2-week drought (T3) and after 5-weeks from the end of drought, *i.e.,* drought recovery (T4). After drought recovery, all above- and belowground functions were collected. Since there was some plant mortality throughout the study, the replication per treatments varied and the observed plant richness was considered at the end (1, 2, 3, 4 species; Table S1) for posterior analyses, in a total of 157 microcosms.

The experiment was carried out in a glasshouse room with corresponding Spring and Summer temperatures from monthly averages (Table S2), from September to January, based on the last 10-year monthly average data from Meteorological Bureau Station 067021 (http://www.bom.gov.au). During the months of plant establishment and as natural light subsidized, 650W LED LumiGrow Pro lights were used to promote plant growth. Microcosms were watered with sterile water every 24–48 hours to maintain soil moisture in the range of 11–14% (corresponding to 60–80% water holding capacity of the soil (WHC)) throughout the experiment.

#### Soil microbial diversity manipulation

Soil was initially collected from the top 0-15 cm at the Yarramundi paddock research site, Western Sydney University in Richmond, NSW, Australia (33°36’34.2”S 150°44’20.9”E). The soil is characterized as loamy sand, with a particle distribution of 81.3% sand, 7.5% silt and 11.2% clay content and a volumetric WHC of 15-20%. It is also characterized by low organic C (1%), total N (0.1%) and pH 5.7. Full soil characteristics can be found in Churchill et al. (2020).

After field collection, soil was sieved with a 5-mm sieve and homogenised and aliquots of approx. 1.8 kg of fresh soil were stored in plastic bags. The bags containing soil were sterilized using gamma radiation (50kGy each) at ANSTO Life Sciences facilities, Sydney, Australia. Gamma radiation was used as it is known to cause minimal change to the physical and chemical properties of soils compared with other methods such as autoclaving (Wolf et al., 1989; Lotrario et al., 1995). A separate portion of soil was kept aside to serve as inoculum and original microbial community characterization. The dilution-to-extinction approach was then used to prepare soil inoculum (Philippot et al., 2013; Delgado-Baquerizo et al., 2020). Six different replicates of inoculum from the unsterilized soil were produced to create a serial dilution to avoid pseudo-replication. The parent inoculum suspensions were prepared by manually mixing 1:4 ratio of soil in sterilized 1X Phosphate buffer saline (PBS) for 5 min. The sediment was then allowed to settle for approx. 2 min and 1:100 serial dilutions were prepared from the suspension using a plate with magnetic stirring to mix the contents homogenously. From these serial dilutions only 3 diversity levels were selected as microbial inoculum, depicting high (HD = 10^0^), moderate (MD = 10^-2^) and low diversity (LD = 10^-6^). The microbial inoculation consisted of a 1:9 ratio of inoculum to soil, leading to soil moisture of approx. 18-20% (corresponding to previously determined WHC). Bags were kept closed with a cotton plug to allow gas exchange and incubated in the dark at room temperature (20°C) for 18 weeks to allow microbial colonization and biomass to recover. The bags were gently mixed every 3 weeks to homogenize microbial growth. Microcosm inoculation and posterior set-up after 18 weeks of incubation was carried within a laminar flow hood and all parts used for the inoculation were sterilized by autoclaving at 121°C. Microcosms consisted of pots (inner diameter = 14 cm, height = 15 cm; volume = 1.9 L) filled with an average of 1.6 kg of soil (dry mass) with the selected microbial diversity levels. After 18 weeks of inoculation, before experimental set-up, and compared to undiluted soil (HD =10^0^), microbial richness of MD was reduced by 9% for bacteria and 14% for fungi and microbial richness of LD was reduced by 52% for bacteria and 73% for fungi.

#### Plant diversity manipulation

Six plant species were initially selected based on species classification into three different functional groups, according to their intrinsic physiological and morphological differences. The species used comprised C4 grasses (*Chloris gayana, Digitaria eriatha*), C3 grasses (*Lolium perenne, Phalaris aquatica*) and legumes (*Biserrula pelecinus, Medicago sativa*). Cultivar information can be found in Churchill et al. (2020). All species were represented in the monocultures (N = 6); and 3 species combination (N = 3) were initially set up as X (*Chloris gayana, Lolium perenne and Biserrula pelecinus*), Y (*Lolium perenne, Phalaris aquatica* and *Medicago sativa*), Z (*Chloris gayana, Biserrula pelecinus, and Medicago sativa*); with all 6 species represented in a final combination (N = 1).

Seedlings of at least 3 cm in height were previously germinated in sterilized vermiculite in growth cabinets (25/15^◦^C day/night; 14/12h day/night) before transplanting them into pots with soil microbial dilutions in the greenhouse. All pots aimed to have six plants, but establishment rate varied. Briefly, monocultures aimed to have six seedlings, three species combinations aimed to have two seedlings per species and six species combination pots aimed to have one seedling per species, to maintain similar evenness. Due to initial low establishment rate, transplant attempts were maintained for the first 3 months and thus it was not possible to account for similar germination times between all species and replicates. Plant diversity (Shannon-Wiener Index; H’) was established at T2 and maintained throughout the duration of the experiment (Figure S1). Seedlings less than 3 cm height as well as dead plants after drought were excluded from plant richness counts and function measurements. In the end, the legume species *Biserrula pelecinus* had the highest mortality and lowest transplant survival and for this reason was removed from further analyses. After plant establishment, the final plant richness achieved was 1, 2, 3 and 4 species from combinations of *Chloris gayana, Digitaria eriatha, Lolium perenne, Phalaris aquatica and Medicago sativa*.

#### Drought manipulation

The drought was applied to all plant-microbial diversity interactions and no plant controls and it consisted half of the initial six replicates being maintained under a reduced watering regime and the other half at a well-watered regime (n=3). For drought replicates, watering was maintained to achieve a soil moisture in the range of 5–7% by weight (30–40% WHC of the soil), whereas well-watered replicates were watered to maintain a moisture content in the range of 11–13% (60-70% WHC). At the end of the drought event (and after sampling), the drought replicates were brought back to well-watered ranges and allowed to recover for five weeks before a final harvest took place.

#### Microbial diversity characterization

Soil was collected for characterization of soil diversity by amplicon sequencing using an Illumina MiSeq platform by Next-Generation Sequencing Facility at the Western Sydney University (Richmond, NSW, Australia). Soil genomic DNA was extracted using the PowerSoil® DNA Isolation kit (Qiagen Inc., USA) according to the manufacturer’s instructions, with modification of the soil weight used (0.50 g) and the initial cell-lysis step, using a FastPrep-24 5G bead beating system (MP Biomedicals, CA, USA) at a speed of 5.5 m s^-1^ for 30 s. Bacterial 16S rRNA and fungal ITS2 region at the start and end of the study were sequenced using 341F/805R and fITS7/ITS4 primer sets, respectively (Delgado-Baquerizo et al., 2016). To quantify the total abundance of bacteria and fungi in the soil at the start and end of the study, the 16S rRNA gene (primer set Eub518-Eub238) and ITS region (primer set ITSIf-5.8s), respectively, were quantified in a Light Cycler 96 Real Time PCR System (Roche) as described in Liu et al. (2019).

Bioinformatic analysis on the 16S rDNA/ITS amplicon reads was performed as previously described. Briefly, pair-end reads were merged using USEARCH (Edgar, 2010) followed by a quality filtering step on the merged reads with expected error (ee) set as 1.0 maximum. High quality reads were then dereplicated and singletons were removed. The zOTUs (zero-radius operational taxonomic units; denoised sequences, 100% sequence identity) were gained by denoising (error-correction) the amplicon reads using unoise3 (Edgar & Flyvbjerg, 2015). Representative sequences of 16S rDNA and ITS zOTUs were annotated against the Silva (Quast et al., 2013) and UNITE (Nilsson et al., 2018) database in QIIME (Caporaso, et al., 2010) using the UCLUST algorithm (Edgar, 2010), respectively. Approximately 15.8M and 17.1 high-quality merged sequences were mapped for 16S rDNA and ITS zOTUs, respectively. Prior to diversity calculation, a normalization procedure was performed at 15,000 and 11,000 sequences *per* sample for 16S rDNA and ITS table, respectively. Alpha diversity indices, including richness, and beta diversity indices including Bray-Curtis matrices were calculated in QIIME.

At the end of the experiment, on average, bacterial communities were dominated by Actinobacteria, followed by Proteobacteria, and Firmicutes; fungal communities were dominated by Ascomycota. We used in this study, microbial richness (number of soil phylotypes) as a metric of soil biodiversity since richness is the simplest and most used metric of biodiversity. The fungal functional groups such as soil mycorrhizal, saprotrophic and plant pathogens fungi were identified using FungalTraits (Põlme et al., 2020). The dilution-to-extinction approach had a significant effect on reducing the soil microbial richness and composition (Figure S2a, S3, S4; Table S3). There was no significant effect of plant richness on total soil bacterial and fungal richness treatment at the end of the experiment, however plant richness had a significant effect on fungal composition (Figure S3). In the microbial richness levels, mycorrhizal fungi decreased on average 76% from HD to MD and LD, saprotrophic fungi decreased 9% and 61% to MD and LD respectively, and plant pathogens decreased 7% and 30% to MD to LD, respectively.

#### Ecosystem Functions

We quantified 16 above- and belowground ecosystem functions regulated by both plant and soil microorganisms and representing direct measures of biotic/abiotic processes as well as process indicators such as nutrient pools (Garland et al., 2021). These functions contribute to climate regulation and support primary production, soil fertility, nutrient cycling, and photosynthesis. We have grouped them into six service categories: soil nutrient storage (total C, N, P), soil inorganic pools (inorganic N, phosphate), OM decomposition (basal respiration, glucose, mineralization, lignin degradation), soil dissolved pools (dissolved organic C (DOC), total dissolved N (TDN)), plant productivity (plant biomass (shoot and root) biomass, height, green canopy cover) and leaf uptake (leaf C, N, P content) (Figure S6). We used substrate induced-respiration measurements as indicators of OM decomposition. Total dissolved N represents the pool of organic and inorganic N. The inclusion of soil dissolved C and N pools provide a more dynamic interpretation of C and N cycles since they represent stocks directly linked to process rates. Similarly, total C, N, P stocks as indicators of nutrient storage are also the result of biological processes and have been found to regulate plant production and diversity in desertification reversal (Qiu et al., 2018; Hu et al., 2021). The 16 variables considered are mostly weakly correlated with each other (Table S4a). Posteriorly, individual functions were grouped into service categories also having weak correlation with each other, further supporting their inclusion in the multifunctionality index and services (Table S4b). All functions and services are also positively correlated to the multifunctionality index (except for a negative correlation between multifunctionality and canopy cover and leaf N, which are non-significant), facilitating the interpretation of the results. At the final harvest, plant height was determined for each individual plant from the ground up and community weighted means were used for 2, 3, and 4-plant species combinations to obtain an average value per pot. Then, shoots were cut at the soil surface, the number of individuals of each species was counted, and shoot per species and total root biomass per pot were determined, by obtaining dry mass after oven-dry at 60°C for 72h. Green canopy cover per pot was obtained using Canopeo (Patrignani & Ochsner, 2015), which provides a measure of “greenness” of the vegetation, and thus acts as a proxy of plant productivity. Leaf and soil C and N were determined on a Vario El Cube CHNS Elemental Analyser (Elementar Australia Pty Lda) and leaf and soil P were determined using analytical Epsilon 3 EDXRF spectrometer (Malvern Panalytical, Leyweg, Almelo, Netherlands) from ground oven-dried samples (soil: 40°C and leaves: 60°C; for 72h) at the end of the experiment. Leaf C, N and P were obtained per species within each plant combination and community weighted means were used. Dissolved organic C and TDN were determined on a total organic C analyser fitted with a total N measurement unit (TOC-L TNM-L, Shimadzu, Sydney, Australia) after a filtered extraction with 0.05M K_2_SO_4_. The availability of inorganic N (sum of ammonium and nitrate) and phosphate (PO_4_^3-^) concentrations in the soil were determined on a SEAL AQ2 Discrete Analyser (SEAL Analytical Inc., USA) after extraction with 2M KCl and 0.5M HCl, respectively. Basal respiration, lignin degradation and glucose mineralization were determined following the MicroResp approach to measure lignin and glucose induced respiration (Campbell et al., 2003). Substrate-induced respiration of glucose and lignin are calculated as respiration in glucose or lignin less the basal respiration. All raw data are made available at Figshare (DOI: 10.6084/m9.figshare.20077217), and sequence data are available at European Nucleotide Archive with accession number PRJEB53575.

### Statistical analysis

#### Multifunctionality index

To obtain a quantitative multifunctionality index three independent multifunctionality approaches were used: (1) the weighted multifunctionality index (2) the multi-threshold multifunctionality index, and (3) multiple individual functions. The weighted ecosystem multifunctionality index was used to prevent the up-weighting of certain aspects of ecosystem functioning or services since some functions were not accounted for between the two data sets, thus providing insight into the average biodiversity effect on services (Manning et al., 2018). It was obtained by first standardizing all individual ecosystem functions between 0 and 1 (rawFunction − min(rawFunction)/(max(rawFunction) − min(rawFunction)); functions were transformed (logarithm or square root) when necessary before standardization), grouped as ecosystem services by averaging individual functions belonging to each group in each data set (three services were considered in the global survey and six services in the microcosm study), and then all services were averaged to account for equal service contribution. Averaged multifunctionality was also included to validate weighted multifunctionality index by simply averaging all individual ecosystem functions equally, without considering ecosystem services grouping. For biodiversity indexes (microbial richness: combination of bacteria and fungi; plant x microbes richness: combination of plant and soil microbes, *i.e.* multitrophic richness) a similar standardization and weighting was applied using the richness of individual groups so that the richness of each group contributed equally to each biodiversity metric. The multithreshold approach was included since it provides insight into the level of performance of multiple functions in response to biodiversity (Byrnes et al., 2014), by estimating the relationship between biodiversity and the number of functions (rate or availability) that simultaneously exceed a critical threshold (>10%, 25%, 50%, 75% and 90% of the maximum observed level of functioning for a given function).

#### Linear mixed modelling

We used a linear mixed effect model approach fitted by restricted maximum likelihood in the microcosm study to test whether soil microbial diversity interacts with plant richness to influence the multiple functions considered. All plant and soil microbial characteristics and all 16 ecosystem functions were assessed for variation among plant and microbial richness and drought treatments after a recovery period. Observed plant richness was considered as a continuous variable (co-variate) due to variation throughout the study. Plant combination (presence/absence of species) was used as a random effect to account for variation from the different plant species established within each combination. Due to low replication at 4-species combination at final harvest, we additionally tested the response of all functions to 3-species diversity, supporting the robustness of our findings. The general response of ecosystem functions and services did not change. When necessary, data were transformed (logarithm or square root) to improve assumptions of normality of errors and homogeneity of variance. Treatment effects were considered statistically significant at *p* < 0.05 and a Tukey HSD post hoc test was used for multiple pairwise comparisons. Linear mixed modelling was performed with JMP v16.0.0 (SAS Institute). A non-metric multidimensional ordination (NMDS) was applied on the matrix of bacterial and fungal composition at the zOTU level using the Bray–Curtis distance and obtained with the package Vegan from R (Oksanen et al., 2017) and a two-way PERMANOVA was applied for treatment effect test.

*Multi-model inference based on information theory.* For the microcosm study, a model-averaging procedure (Burnham & Anderson, 2002) was employed for multifunctionality, ecosystem services and individual functions separately, based on minimizing the corrected Akaike information criterion (AICc) to evaluate the % of importance of the predictors under consideration, namely plant richness and composition and microbial richness and drought as drivers of multifunctionality. We included plant composition in the microcosm study analysis to account for the lack of similar germination times between all species and replicates which can contribute to variation in final plant composition rates. All predictors and response variables were standardized before analyses to interpret parameter estimates on a comparable scale. This method is similar to a variance partition analysis because of previous standardization of predictors (Gross et al., 2017; Le Bagousse-Pinguet et al., 2019; Le Provost et al., 2020; Le Bagousse-Pinguet et al., 2021). We removed microbial composition as well as plant and soil microbial richness combined as predictors due to high correlation (Table S4c). Microbial richness was obtained from the mean of standardized bacterial and fungal diversity, so each group contribution is accounted equally. Plant species composition was estimated from shoot biomass of each species as % Composition species a = 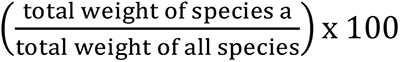 and was obtained from the first axis of a principal component analysis (PCA; Figure S5a). Model residuals were inspected to ensure for constant variance and normality. The importance of predictors was expressed as the percentage of variance they explain, based on the comparison between the absolute values of their standardized regression coefficients and the sum of all standardized regression coefficients from all predictors in the models. R^2^ values presented express total variances corresponding to model adj. R² obtained from parameter estimates averaged across all models.

A similar approach was applied for the global survey where aridity index was included and environmental properties (distance from the equator, plant cover, soil pH, % clay, soil C and mean annual temperature) corresponded to a standardized first axis of a PCA analysis obtained from the multiple properties (Figure S5b). We did not include plant composition for the global grassland survey because obtaining absolute abundance of plant species at the global scale was not possible. We excluded the multitrophic richness as predictors in both datasets due to high correlation (Table S4c). Multi-model analyses were carried out using SAM 4.0 (Rangel et al., 2010).

#### Regression analysis and spearman correlations

In both global and microcosm data sets, to test the relationship between the standardized individual richness groups (plant richness, bacterial richness, fungal richness, mycorrhizal, saprotrophic and plant pathogens fungal richness) and multitrophic richness (joint effects of plant and soil microbial richness) with ecosystem multifunctionality (weighted and multi-threshold multifunctionality), we used best fitted regressions (linear, quadratic) Spearman correlations between richness groups and weighted multifunctionality, ecosystem services and individual functions were also conducted.

#### Partial correlations

For the microcosm study, to test for the influence of community composition and abundance in biodiversity–multifunctionality relationships, we conducted partial correlation analysis between plant and soil biodiversity and weighted multifunctionality, accounting for plant abundance (biomass) and microbial abundance (quantitative PCR data) and plant composition (main axis of a PCA analysis) and microbial composition (main axis of a NMDS analysis). Regression analysis, Spearman correlations and partial correlations were performed with JMP v16.0.0 (SAS Institute).

#### Structural equation modelling (SEM)

We used SEM to evaluate the direct link between plant and soil biodiversity and weighted multifunctionality in the global survey, after accounting for multiple key environmental factors such as spatial influence (distance from equator), climate (mean annual temperature and aridity), plant (richness and cover) and soil (soil pH, total C content and percentage of clay) attributes (Table S5a for standardized direct effects considered in the a priori model). In contrast to regression analysis, SEM offers the ability to separate multiple pathways of influence and view them as parts of a system and is therefore useful for investigating the complex relationships among predictors (Delgado-Baquerizo et al., 2016). Full details of approach followed can be found in (Delgado-Baquerizo et al., 2020). We also used SEM to evaluate the effects of plant, bacterial and fungal richness on above- and belowground functions after accounting for influences of drought and plant composition in the microcosm study. Drought was included as a categorical variable with two levels: 1 (well-watered) and 0 (drought). All remaining variables were included as independent observable variables, with plant composition consisting of the main axis of a PCA analysis obtained from the final % of composition of each plant species. We considered that plant richness influenced final plant composition by the end of the experiment since plant composition was dependent and thus derived from the experimentally manipulated plant richness. Due to the duration of microcosm study, we also considered that by the end, there would be an influence of microbial richness on the plant composition since this metric was derived from final shoot biomass. All standardized direct effects considered in the a priori model can be found in Table S5b. Model overall goodness-of-fit was tested using the Chi-square test (*χ*2; the model has a good fit when 0 ≤ *χ*2 ≤ 2 and 0.05 < *p* ≤ 1.00) and the root mean square error of approximation (RMSEA; the model has a good fit when 0 ≤ RMSEA ≤ 0.05 and 0.10 < *p* ≤ 1.00; (Schermelleh-Engel et al., 2003). Additionally, because some variables in the microcosm study resulted in residuals that were not normally distributed, the model fit was confirmed using the Bollen-Stine bootstrap test (the model has a good fit when 0.10 < bootstrap *p* ≤ 1.00). Structural equation modelling was performed using AMOS 27.0, IBM SPSS.

## Data availability

Metadata and samples ID are freely available in Figshare (https://doi.org/10.6084/m9.figshare.20077217).

## Authors’ contributions

C.S.C.M., M.D-B., F.T.M. P.B.R and B.K.S conceived and designed the original ideas of this study. The field dataset was supervised by F.T.M., M.D-B., and B.K.S. The microcosm experiment was conducted by C.S.C.M. with the assistance of R.H.J. Q-PCR analysis was carried out by H.L. Bioinformatics analyses were performed by J-T.W. Statistical analyses were performed by C.S.C.M in consultation with M.D.-B. The first draft of the paper was written by C.S.C.M., with subsequent revisions by M.D.-B., F.T.M., P.B.R and B.K.S. All authors contributed to the final draft.

## Acknowledgements

The project was funded by Australia Research Council (DP170104634 and DP210102081). M.D-B. acknowledges support from the Spanish Ministry of Science and Innovation for the I+D+i project PID2020-115813RA-I00 funded by MCIN/AEI/10.13039/501100011033. M.D-B. is also supported by a project of the Fondo Europeo de Desarrollo Regional (FEDER) and the Consejería de Transformación Económica, Industria, Conocimiento y Universidades of the Junta de Andalucía (FEDER Andalucía 2014-2020 Objetivo temático “01 – Refuerzo de la investigación, el desarrollo tecnológico y la innovación”) associated with the research project P20_00879 (ANDABIOMA). F.T.M. acknowledges support from the European Research Council (ERC Grant agreement 647038 [BIODESERT]) and Generalitat Valenciana (CIDEGENT/2018/041). P.B.R. acknowledges support by the National Science Foundation, Biological Integration Institutes grant NSF-DBI-2021898. For the experimental study (microcosm), authors thank the technical support in sample collections and laboratory analysis, particularly B. Batista, P. Singh, P. Matta and D. Wojtalewicz.

## References

Bardgett, R. D., Bullock, J. M., Lavorel, S., Manning, P., Schaffner, U., Ostle, N., … Shi, H. (2021). Combatting global grassland degradation. Nature Reviews Earth & Environment. doi:10.1038/s43017-021-00207-2

Bardgett, R. D., & van der Putten, W. H. (2014). Belowground biodiversity and ecosystem functioning. Nature, 515(7528), 505–511. doi:10.1038/nature13855

Bardgett, R. D., & Wardle, D. A. (2010). Aboveground-belowground linkages: biotic interactions, ecosystem processes, and global change: Oxford University Press.

Barry, K. E., Mommer, L., van Ruijven, J., Wirth, C., Wright, A. J., Bai, Y., … Weigelt, A. (2019). The Future of Complementarity: Disentangling Causes from Consequences. Trends in Ecology & Evolution, 34(2), 167–180. doi:https://doi.org/10.1016/j.tree.2018.10.013

Begum, N., Qin, C., Ahanger, M. A., Raza, S., Khan, M. I., Ashraf, M., … Zhang, L. (2019). Role of Arbuscular Mycorrhizal Fungi in Plant Growth Regulation: Implications in Abiotic Stress Tolerance. Frontiers in Plant Science, 10(1068). doi:10.3389/fpls.2019.01068

Bengtsson, J., Bullock, J. M., Egoh, B., Everson, C., Everson, T., O’Connor, T., … Lindborg, R. (2019). Grasslands—more important for ecosystem services than you might think. Ecosphere, 10(2), e02582. doi: https://doi.org/10.1002/ecs2.2582

Berdugo, M., Delgado-Baquerizo, M., Soliveres, S., Hernández-Clemente, R., Zhao, Y., Gaitán, J. J., … Maestre, F.T. (2020). Global ecosystem thresholds driven by aridity. Science, 367(6479), 787–790. doi:doi: 10.1126/science.aay5958

Bond, W. J. (2016). Ancient grasslands at risk. Science, 351(6269), 120–122. doi:doi:10.1126/science.aad5132

Bowker, M. A., Rengifo-Faiffer, M. C., Antoninka, A. J., Grover, H. S., Coe, K. K., Fisher, K., … Stark, L.R.. (2021). Community composition influences ecosystem resistance and production more than species richness or intraspecific diversity. Oikos, 130(8), 1399–1410. doi:https://doi.org/10.1111/oik.08473

Burnham, K. P., & Anderson, D. R. (2002). Model selection and multimodel inference. A practical information-theoretical approach. New York: Springer-Verlag.

Burrell, A. L., Evans, J. P., & De Kauwe, M. G. (2020). Anthropogenic climate change has driven over 5 million km2 of drylands towards desertification. Nature Communications, 11(1), 3853. doi:10.1038/s41467-020-17710-7

Byrnes, J. E. K., Gamfeldt, L., Isbell, F., Lefcheck, J. S., Griffin, J. N., Hector, A., … Emmett Duffy, J. (2014). Investigating the relationship between biodiversity and ecosystem multifunctionality: challenges and solutions. Methods in Ecology and Evolution, 5(2), 111–124. doi:https://doi.org/10.1111/2041-210X.12143

Cameron, E. K., Martins, I. S., Lavelle, P., Mathieu, J., Tedersoo, L., Bahram, M., … Eisenhauer, N. (2019). Global mismatches in aboveground and belowground biodiversity. Conservation Biology, 33(5), 1187–1192. doi:https://doi.org/10.1111/cobi.13311

Campbell, C. D., Chapman, S. J., Cameron, C. M., Davidson, M. S., & Potts, J. M. (2003). A rapid microtiter plate method to measure carbon dioxide evolved from carbon substrate amendments so as to determine the physiological profiles of soil microbial communities by using whole soil. Applied and Environmental Microbiology, 69(6), 3593–3599. doi:10.1128/aem.69.6.3593-3599.2003

Caporaso, J. G., Kuczynski, J., Stombaugh, J., Bittinger, K., Bushman, F. D., Costello, E. K., … Knight, R. (2010). QIIME allows analysis of high-throughput community sequencing data. Nature Methods, 7(5), 335–336. doi:10.1038/nmeth.f.303

Cardinale, B. J., Matulich, K. L., Hooper, D. U., Byrnes, J. E., Duffy, E., Gamfeldt, L., … Gonzalez, A. (2011). The functional role of producer diversity in ecosystems. American Journal of Botany, 98(3), 572–592. doi:https://doi.org/10.3732/ajb.1000364

Carrick, P. J., & Forsythe, K. J. (2020). The species composition—ecosystem function relationship: A global meta-analysis using data from intact and recovering ecosystems. PLoS ONE, 15(7), e0236550. doi:10.1371/journal.pone.0236550

Chen, M., Arato, M., Borghi, L., Nouri, E., & Reinhardt, D. (2018). Beneficial Services of Arbuscular Mycorrhizal Fungi – From Ecology to Application. Frontiers in Plant Science, 9(1270). doi:10.3389/fpls.2018.01270

Churchill, A. C., Zhang, H., Fuller, K. J., Amiji, B., Anderson, I. C., Barton, C. V. M., … Power, S.A.. (2020). Pastures and Climate Extremes: Impacts of warming and drought on the productivity and resilience of key pasture species in a field experiment. bioRxiv, 2020.2012.2021.423155. doi:10.1101/2020.12.21.423155

Crawford, J. W., Deacon, L., Grinev, D., Harris, J. A., Ritz, K., Singh, B. K., Young, I. (2012). Microbial diversity affects self-organization of the soil-microbe system with consequences for function. Journal of the Royal Society Interface 9, 1302–1310.

de Vries, F. T., Griffiths, R. I., Bailey, M., Craig, H., Girlanda, M., Gweon, H. S., … Bardgett, R.D. (2018). Soil bacterial networks are less stable under drought than fungal networks. Nature Communications, 9(1), 3033. doi:10.1038/s41467-018-05516-7

Delgado-Baquerizo, M., Maestre, F. T., Gallardo, A., Bowker, M. A., Wallenstein, M. D., Quero, J. L., … Zaady, E. (2013). Decoupling of soil nutrient cycles as a function of aridity in global drylands. Nature, 502(7473), 672–676. doi:10.1038/nature12670

Delgado-Baquerizo, M., Maestre, F. T., Reich, P. B., Jeffries, T. C., Gaitan, J. J., Encinar, D., … Singh, B.K. (2016). Microbial diversity drives multifunctionality in terrestrial ecosystems. Nature Communications, 7, 10541. doi:10.1038/ncomms10541

Delgado-Baquerizo, M., Bardgett, R. D., Vitousek, P. M., Maestre, F. T., Williams, M. A., Eldridge, D. J., … Fierer, N. (2019). Changes in belowground biodiversity during ecosystem development. Proceedings of the National Academy of Sciences, 116(14), 6891–6896. doi:doi:10.1073/pnas.1818400116

Delgado-Baquerizo, M., Reich, P. B., Trivedi, C., Eldridge, D. J., Abades, S., Alfaro, F. D., … Singh, B.K.. (2020). Multiple elements of soil biodiversity drive ecosystem functions across biomes. Nature Ecology & Evolution, 4(2), 210–220. doi:10.1038/s41559-019-1084-y

Eastwood, D. C., Floudas, D., Binder, M., Majcherczyk, A., Schneider, P., Aerts, A., … Watkinson, S.C. (2011). The Plant Cell Wall&#x2013;Decomposing Machinery Underlies the Functional Diversity of Forest Fungi. Science, 333(6043), 762–765. doi:doi:10.1126/science.1205411

Edgar, R. C. (2010). Search and clustering orders of magnitude faster than BLAST. Bioinformatics, 26(19), 2460–2461.

Edgar, R. C., & Flyvbjerg, H. (2015). Error filtering, pair assembly and error correction for next-generation sequencing reads. Bioinformatics, 31(21), 3476–3482.

Garland, G., Banerjee, S., Edlinger, A., Miranda Oliveira, E., Herzog, C., Wittwer, R., … van der Heijden, M. G. A. (2021). A closer look at the functions behind ecosystem multifunctionality: A review. Journal of Ecology, 109(2), 600–613. doi:https://doi.org/10.1111/1365-2745.13511

Gessner, M. O., Swan, C. M., Dang, C. K., McKie, B. G., Bardgett, R. D., Wall, D. H., & Hättenschwiler, S. (2010). Diversity meets decomposition. Trends in Ecology & Evolution, 25(6), 372–380. doi:https://doi.org/10.1016/j.tree.2010.01.010

Gross, N., Bagousse-Pinguet, Y. L., Liancourt, P., Berdugo, M., Gotelli, N. J., & Maestre, F. T. (2017). Functional trait diversity maximizes ecosystem multifunctionality. Nature Ecology & Evolution, 1(5), 0132. doi:10.1038/s41559-017-0132

Habel, J. C., Dengler, J., Janišová, M., Török, P., Wellstein, C., & Wiezik, M. (2013). European grassland ecosystems: threatened hotspots of biodiversity. Biodiversity and Conservation, 22(10), 2131–2138. doi:10.1007/s10531-013-0537-x

Hooper, D. U., Bignell, D. E., Brown, V. K., Brussard, L., Dangerfield, J. M., Wall, D. H., … Wolters, V. (2000). Interactions between Aboveground and Belowground Biodiversity in Terrestrial Ecosystems: Patterns, Mechanisms, and Feedbacks: We assess the evidence for correlation between aboveground and belowground diversity and conclude that a variety of mechanisms could lead to positive, negative, or no relationship— depending on the strength and type of interactions among species. BioScience, 50(12), 1049–1061. doi: 10.1641/0006-3568(2000)050[1049:Ibaabb]2.0.Co;2

Hooper, D. U., & Vitousek, P. M. (1998). Effects of Plant Composition and Diversity on Nutrient Cycling. Ecological Monographs, 68(1), 121–149. doi:10.2307/2657146

Hu, W., Ran, J., Dong, L., Du, Q., Ji, M., Yao, S., … Deng, J. (2021). Aridity-driven shift in biodiversity–soil multifunctionality relationships. Nature Communications, 12(1), 5350. doi:10.1038/s41467-021-25641-0

Huang, J., Yu, H., Guan, X., Wang, G., & Guo, R. (2016). Accelerated dryland expansion under climate change. Nature Climate Change, 6(2), 166–171. doi:10.1038/nclimate2837

IPCC. (2013). Summary for Policymakers. In T. F. Stocker, D., Qin, G.-K., Plattner, M. Tignor, S. K. Allen, J. Boschung, A. Nauels, Y., Xia, V. Bex, & P. M. Midgley (Eds.), Climate Change 2013: The Physical Science Basis. Contribution of Working Group I to the Fifth Assessment Report of the Intergovernmental Panel on Climate Change (pp. 1–30). Cambridge, United Kingdom and New York, NY, USA: Cambridge University Press.

Isbell, F., Calcagno, V., Hector, A., Connolly, J., Harpole, W. S., Reich, P. B., … Loreau, M. (2011). High plant diversity is needed to maintain ecosystem services. Nature, 477(7363), 199–202. doi:10.1038/nature10282

Jing, X., Sanders, N. J., Shi, Y., Chu, H., Classen, A. T., Zhao, K., … He, J.-S. (2015). The links between ecosystem multifunctionality and above- and belowground biodiversity are mediated by climate. Nature Communications, 6(1), 8159. doi:10.1038/ncomms9159

Laliberté, E., Zemunik, G., & Turner, B. L. (2014). Environmental filtering explains variation in plant diversity along resource gradients. Science, 345(6204), 1602–1605. doi:10.1126/science.1256330

Le Bagousse-Pinguet, Y., Gross, N., Saiz, H., Maestre, F. T., Ruiz, S., Dacal, M., … García-Palacios, P. (2021). Functional rarity and evenness are key facets of biodiversity to boost multifunctionality. Proceedings of the National Academy of Sciences, 118(7), e2019355118. doi:10.1073/pnas.2019355118

Le Bagousse-Pinguet, Y., Soliveres, S., Gross, N., Torices, R., Berdugo, M., & Maestre, F. T. (2019). Phylogenetic, functional, and taxonomic richness have both positive and negative effects on ecosystem multifunctionality. Proceedings of the National Academy of Sciences, 116(17), 8419–8424. doi:10.1073/pnas.1815727116

Le Provost, G., Badenhausser, I., Le Bagousse-Pinguet, Y., Clough, Y., Henckel, L., Violle, C., … Gross, N. (2020). Land-use history impacts functional diversity across multiple trophic groups. Proceedings of the National Academy of Sciences, 117(3), 1573–1579. doi:10.1073/pnas.1910023117

Li, J., Meng, B., Chai, H., Yang, X., Song, W., Li, S., … Sun, W. (2019). Arbuscular Mycorrhizal Fungi Alleviate Drought Stress in C3 (Leymus chinensis) and C4 (Hemarthria altissima) Grasses via Altering Antioxidant Enzyme Activities and Photosynthesis. Frontiers in Plant Science, 10(499). doi:10.3389/fpls.2019.00499

Liu, H., Khan, M. Y., Carvalhais, L. C., Delgado-Baquerizo, M., Yan, L., Crawford, M., … Schenk, P. M.. (2019). Soil amendments with ethylene precursor alleviate negative impacts of salinity on soil microbial properties and productivity. Scientific Reports, 9(1), 6892. doi:10.1038/s41598-019-43305-4

Lotrario, J. B., Stuart, B. J., Lam, T., Arands, R. R., O’Connor, O. A., & Kosson, D. S. (1995). Effects of sterilization methods on the physical characteristics of soil: Implications for sorption isotherm analyses. Bulletin of Environmental Contamination and Toxicology, 54(5), 668–675. doi:10.1007/BF00206097

Lu, M., & Hedin, L. O. (2019). Global plant–symbiont organization and emergence of biogeochemical cycles resolved by evolution-based trait modelling. Nature Ecology & Evolution, 3(2), 239–250. doi:10.1038/s41559-018-0759-0

Ma, Y., Dias, M. C., & Freitas, H. (2020). Drought and Salinity Stress Responses and Microbe–Induced Tolerance in Plants. Frontiers in Plant Science, 11(1750). doi:10.3389/fpls.2020.591911

Maestre, F. T., Quero, J. L., Gotelli, N. J., Escudero, A., Ochoa, V., Delgado-Baquerizo, M., … Zaady, E. (2012). Plant species richness and ecosystem multifunctionality in global drylands. Science, 335(6065), 214–218. doi:10.1126/science.1215442

Manning, P., van der Plas, F., Soliveres, S., Allan, E., Maestre, F. T., Mace, G., … Fischer, M. (2018). Redefining ecosystem multifunctionality. Nature Ecology & Evolution, 2(3), 427–436. doi:10.1038/s41559-017-0461-7

Manzoni, S., Schimel, J. P., & Porporato, A. (2012). Responses of soil microbial communities to water stress: results from a meta-analysis. Ecology, 93(4), 930–938. doi:10.1890/11-0026.1

Nguyen, N. H., Song, Z., Bates, S. T., Branco, S., Tedersoo, L., Menke, J., … Kennedy, P. G.. (2016). FUNGuild: An open annotation tool for parsing fungal community datasets by ecological guild. Fungal Ecology, 20, 241–248. doi:https://doi.org/10.1016/j.funeco.2015.06.006

Nilsson, R. H., Larsson, K.-H., Taylor, A. F S., Bengtsson-Palme, J., Jeppesen, T. S., Schigel, D., … Abarenkov, K. (2018). The UNITE database for molecular identification of fungi: handling dark taxa and parallel taxonomic classifications. Nucleic acids research, 47(D1), D259–D264. doi:10.1093/nar/gky1022

O’Mara, F. P. (2012). The role of grasslands in food security and climate change. Annals of Botany, 110(6), 1263–1270. doi:10.1093/aob/mcs209

Ochoa-Hueso, R., Eldridge, D. J., Delgado-Baquerizo, M., Soliveres, S., Bowker, M. A., Gross, N., … Maestre, F. T. (2018). Soil fungal abundance and plant functional traits drive fertile island formation in global drylands. Journal of Ecology, 106(1), 242–253. doi:https://doi.org/10.1111/1365-2745.12871

Oleńska, E., Małek, W., Wójcik, M., Swiecicka, I., Thijs, S., & Vangronsveld, J. (2020). Beneficial features of plant growth-promoting rhizobacteria for improving plant growth and health in challenging conditions: A methodical review. Science of The Total Environment, 743, 140682. doi:https://doi.org/10.1016/j.scitotenv.2020.140682

Oksanen, J., Blanchet, G. F., Friendly, M., Kindt, R., Legendre, P., McGlinn, D., … Wagner, H. (2017). *vegan: Community Ecology Package. R package version 2.4-3*. retrieved from https://CRAN.R-project.org/package=vegan

Orwin, K. H., Kirschbaum, M. U. F., St John, M. G., & Dickie, I. A. (2011). Organic nutrient uptake by mycorrhizal fungi enhances ecosystem carbon storage: a model-based assessment. Ecology Letters, 14(5), 493–502. doi:https://doi.org/10.1111/j.1461-0248.2011.01611.x

Patrignani, A., & Ochsner, T. (2015). Canopeo: A Powerful New Tool for Measuring Fractional Green Canopy Cover. Agronomy Journal, 107, 2312–2320.

Philippot, L., Spor, A., Hénault, C., Bru, D., Bizouard, F., Jones, C. M., … Maron, P.-A. (2013). Loss in microbial diversity affects nitrogen cycling in soil. The ISME Journal, 7(8), 1609–1619. doi:10.1038/ismej.2013.34

Põlme, S., Abarenkov, K., Henrik Nilsson, R., Lindahl, B. D., Clemmensen, K. E., Kauserud, H., … Tedersoo, L. (2020). FungalTraits: a user-friendly traits database of fungi and fungus-like stramenopiles. Fungal Diversity, 105(1), 1–16. doi:10.1007/s13225-020-00466-2

Qiu, K., Xie, Y., Xu, D., & Pott, R. (2018). Ecosystem functions including soil organic carbon, total nitrogen and available potassium are crucial for vegetation recovery. Scientific Reports, 8(1), 7607. doi:10.1038/s41598-018-25875-x

Quast, C., Pruesse, E., Yilmaz, P., Gerken, J., Schweer, T., Yarza, P., … Glöckner, F.O. (2013). The SILVA ribosomal RNA gene database project: improved data processing and web-based tools. Nucleic acids research, 41(Database issue), D590–D596. doi:10.1093/nar/gks1219

Rangel, T. F., Diniz-Filho, J. A. F., & Bini, L. M. (2010). SAM: a comprehensive application for Spatial Analysis in Macroecology. Ecography, 33(1), 46–50. doi:10.1111/j.1600-0587.2009.06299.x

Reich, P. B., Tilman, D., Isbell, F., Mueller, K., Hobbie, S. E., Flynn, D. F. B., & Eisenhauer, N. (2012). Impacts of Biodiversity Loss Escalate Through Time as Redundancy Fades. Science, 336(6081), 589–592. doi:10.1126/science.1217909

Schermelleh-Engel, K., Moosbrugger, H., & Müller, H. (2003). Evaluating the Fit of Structural Equation Models: Tests of Significance and Descriptive Goodness-of-Fit Measures. Methods of Psychological Research, 8(2), 23–74.

Schimel, J. P. (2018). Life in Dry Soils: Effects of Drought on Soil Microbial Communities and Processes. Annual Review of Ecology, Evolution, and Systematics, 49(1), 409–432. doi:10.1146/annurev-ecolsys-110617-062614

Sexton, A. N., & Emery, S. M. (2020). Grassland restorations improve pollinator communities: a meta-analysis. Journal of Insect Conservation, 24(4), 719–726. doi:10.1007/s10841-020-00247-x

Sherwood, S., & Fu, Q. (2014). A Drier Future? Science, 343(6172), 737–739. doi:10.1126/science.1247620

Siciliano, S. D., Palmer, A. S., Winsley, T., Lamb, E., Bissett, A., Brown, M. V., … Snape, I. (2014). Soil fertility is associated with fungal and bacterial richness, whereas pH is associated with community composition in polar soil microbial communities. Soil Biology and Biochemistry, 78, 10–20. doi:https://doi.org/10.1016/j.soilbio.2014.07.005

Soliveres, S., van der Plas, F., Manning, P., Prati, D., Gossner, M. M., Renner, S. C., … Allan, E. (2016). Biodiversity at multiple trophic levels is needed for ecosystem multifunctionality. Nature, 536(7617), 456–459. doi:10.1038/nature19092

Suttie, J. M., Reynolds, S. G., & Batello, C. (2005). Grasslands of the world. FAO.

Tedersoo, L., Bahram, M., & Zobel, M. (2020). How mycorrhizal associations drive plant population and community biology. Science, 367(6480), eaba1223. doi:10.1126/science.aba1223

Tilman, D., Isbell, F., & Cowles, J. M. (2014). Biodiversity and Ecosystem Functioning. Annual Review of Ecology, Evolution, and Systematics, 45(1), 471–493. doi:10.1146/annurev-ecolsys-120213-091917

Tilman, D., Wedin, D., & Knops, J. (1996). Productivity and sustainability influenced by biodiversity in grassland ecosystems. Nature, 379(6567), 718–720. doi:10.1038/379718a0

Trivedi C, Delgado-Baquerizo M, Hamonts K, Lai K, Reich PB, Singh BK. (2019). Losses in microbial functional diversity reduce the rate of key soil processes. Soil Biology and Biochemistry.135, 267–74.

Van Der Heijden, M. G. A., Bardgett, R. D., & Van Straalen, N. M. (2008). The unseen majority: soil microbes as drivers of plant diversity and productivity in terrestrial ecosystems. Ecology Letters, 11(3), 296–310. doi:10.1111/j.1461-0248.2007.01139.x

Wagg, C., Bender, S. F., Widmer, F., & van der Heijden, M. G. A. (2014). Soil biodiversity and soil community composition determine ecosystem multifunctionality. Proceedings of the National Academy of Sciences, 111(14), 5266–5270. doi:10.1073/pnas.1320054111

Wagg, C., Schlaeppi, K., Banerjee, S., Kuramae, E. E., & van der Heijden, M. G. A. (2019). Fungal-bacterial diversity and microbiome complexity predict ecosystem functioning. Nature Communications, 10(1), 4841. doi:10.1038/s41467-019-12798-y

Wall, D. H., Nielsen, U. N., & Six, J. (2015). Soil biodiversity and human health. Nature, 528(7580), 69–76. doi:10.1038/nature15744

Wardle, D. A., Bardgett, R. D., Klironomos, J. N., Setälä, H., van der Putten, W. H., & Wall, D. H. (2004). Ecological Linkages Between Aboveground and Belowground Biota. Science, 304(5677), 1629–1633. doi:10.1126/science.1094875

Wolf, D. C., Dao, T. H., Scott, H. D., & Lavy, T. L. (1989). Influence of Sterilization Methods on Selected Soil Microbiological, Physical, and Chemical Properties. Journal of Environmental Quality, 18(1), 39–44. https://doi.org/10.2134/jeq1989.00472425001800010007x

Yang, G., Ryo, M., Roy, J., Hempel, S., & Rillig, M. C. (2021). Plant and soil biodiversity have non-substitutable stabilising effects on biomass production. Ecology Letters, n/a(n/a). doi:https://doi.org/10.1111/ele.13769

Yang, Y., Tilman, D., Furey, G., & Lehman, C. (2019). Soil carbon sequestration accelerated by restoration of grassland biodiversity. Nature Communications, 10(1), 718. doi:10.1038/s41467-019-08636-w

Zhao, M., Xue, K., Wang, F., Liu, S., Bai, S., Sun, B., … Yang, Y. (2014). Microbial mediation of biogeochemical cycles revealed by simulation of global changes with soil transplant and cropping. The ISME Journal, 8(10), 2045–2055. doi:10.1038/ismej.2014.46

Zhou, Z., Wang, C., & Luo, Y. (2020). Meta-analysis of the impacts of global change factors on soil microbial diversity and functionality. Nature Communications, 11(1), 3072. doi:10.1038/s41467-020-16881-7

